# 3’tRNA^Asp(GTC)^-derived fragment links inflammation to post-transcriptional reprogramming in chondrocytes during osteoarthritis

**DOI:** 10.64898/2026.01.23.700816

**Authors:** Francesco Alabiso, Aleksandra Seragnoli Chystyakova, Cristina Cosentino, Irene Bissoli, Veronique Menoud, Ilaria Bedei, Merissa Olmer, Veronica Panichi, Isabella Rusciano, Paolo Dolzani, Carla Renata Arciola, Stefano Ratti, Martin K. Lotz, Rosa Maria Borzì, Romano Regazzi, Flavio Flamigni, Silvia Cetrullo, Stefania D’Adamo

## Abstract

**Objective:** Transfer RNA-derived fragments (tRFs) belong to an emerging class of small non-coding RNAs that dynamically respond to metabolic stressors and drive different pathological processes, yet their role in osteoarthritis (OA) remains poorly explored. We aimed to define the tRF landscape in OA and investigate the function of 3’tRF^Asp(GTC)^ in chondrocyte stress adaptation and translational control.

**Methods:** *Ex vivo*, cartilage specimens from OA patients (n=6) and healthy donors (n=7) were analyzed by small RNA sequencing to define disease-associated tRF signatures. *In vitro*, primary chondrocytes derived from OA patients were treated with lipopolysaccharide (LPS) to mimic inflammatory environment of OA, used for small RNA sequencing (n=3) and validation analysis (n=6). Functional studies in C28/I2 chondrocytes included antisense oligonucleotide-mediated 3’tRF^Asp(GTC)^ inhibition, AGO2-RNA immunoprecipitation (RIP), polysome profiling, stress granule (SG) immunofluorescence, and differential protein analysis. Computational target prediction and pathway enrichment were used to explore tRF-mediated regulatory networks.

**Results:** Both OA cartilage and LPS-treated chondrocytes displayed upregulation of 3’tRF^Asp(GTC)^ and 5’tRF^Glu(CTC)^, indicating a shared inflammatory tRF signature. Predicted targets of upregulated tRFs were enriched in stress-adaptive, proteostasis, and translational control pathways, whereas downregulated tRFs modulated mitochondrial processes. Silencing 3’tRF^Asp(GTC)^ inhibited LPS-induced *COX2* and *MMP13* expression, prevented ER stress, and blocked SG assembly. RIP confirmed selective recruitment of 3’tRF^Asp(GTC)^ into AGO2 complexes. Polysome profiling revealed association with 40S ribosomal subunit, mediating translational arrest and influencing selective mRNA expression.

**Conclusion:** 3’tRF^Asp(GTC)^ emerges as a regulator linking inflammation to translational control and SG dynamics in OA. tRFs thus could represent novel therapeutic targets in OA disease.

## Introduction

Osteoarthritis (OA) is characterized by cartilage degradation and low-grade inflammation. Beyond aging, metabolic abnormalities such as obesity and type II diabetes contribute to OA pathogenesis independently of mechanical loading^1, 2^. In this context, metabolic inflammation promotes innate immune activation within the joint, while circulating inflammatory mediators and metabolic endotoxins, including lipopolysaccharides (LPS), further drive cartilage catabolism through Toll-like receptor 4 (TLR4) signalling^3–5^. OA shares aberrant molecular features with other degenerative diseases, including alterations of non-coding RNAs (ncRNA) and stress granule (SG) assembly, both involved in the post-transcriptional regulation of gene expression^6–8^. However, the mediators underlying these processes remain largely unclear.

Transfer RNAs (tRNAs) are housekeeping ncRNAs bearing the correct amino acid during translation. Enzyme-mediated tRNA cleavage produces tRNA-derived fragments (tRFs), a novel class of highly dynamic ncRNAs that exert their functions by binding with a variety of molecular partners^9, 10^. tRFs are classified as 5’, 3’ or internal fragments based on the cleavage site, with tRNA-halves (or tiRNAs, ∼30 nt) generated by anticodon-loop cleavage distinguished from other tRFs of varying lengths^10^. In archaea and yeast, tRFs have been shown to directly influence translation, interacting with ribosomes^9, 11^. Consistently, tRFs have been described to be involved in translation modulation by regulating SG assembly in U2OS cells^12^. The formation of ribonucleoprotein (RNP) complexes, including SGs, is one of the processes engaged by cells to counteract metabolic stress. It is strictly linked to the unfolded protein response (UPR) and mediates translational reprogramming^13, 14^. These processes suppress bulk protein synthesis while favouring specific stress-response transcripts. Untranslated mRNAs accumulate in SGs, whose assembly depends on RNA-binding proteins (RBPs) such as TIA-1, HuR, and G3BPs, enabling dynamic storage or selective translation^15^. Despite evidence linking tRFs to translational regulation, their role in remodelling translation during OA pathogenesis remains unexplored.

This study provides an unbiased characterization of the tRF signature distinguishing OA patients from healthy individuals and integrates these findings into a controlled *in vitro* model. Using LPS to recapitulate the pro-inflammatory milieu of metabolic OA, we interrogated the interplay between tRF biogenesis, translational status, and SG dynamics, with the aim of defining how disease-associated tRFs orchestrate translational control and inflammatory stress adaptation in chondrocytes.

## Methods

### Cartilage donors

Normal human knee cartilage tissues were procured by tissue banks from seven donors without history of joint disease or trauma. Full-thickness cartilage was harvested for RNA isolation from the medial and lateral femoral condyles. OA-affected cartilage was harvested from six donors during knee replacement surgery. Cartilage specimens were stored at −80°C immediately after collection. These human tissues were obtained with approval from the Scripps Human Subjects Committee.

### Primary Chondrocyte Isolation

Human articular chondrocytes were isolated from femoral condyles and tibial plateaus of OA patients undergoing total knee arthroplasty at the Rizzoli Orthopaedic Institute. All donors provided written informed consent, and the study was approved by the local ethics committee (CE AVEC 209/2022/Sper/IOR). Isolation procedures were performed as previously described^5^. Primary chondrocytes were used for small RNA sequencing and Real-Time qPCR validation.

### C28/I2 Cell Culture

The human immortalized chondrocyte line C28/I2 was cultured in DMEM (Gibco) supplemented with 10% FBS (Gibco) and 1% penicillin/streptomycin (Sigma) and used for antisense oligonucleotide (ASO) experiments, SG analysis, and immunoprecipitation/polysome profiling assays.

### In Vitro OA Model and Treatments

Primary and C28/I2 chondrocytes were plated at high density (2.5 × 10⁵ cells/well in 12-well plates) 48–72 h before treatment. Inflammatory stress was induced with LPS (10 µg/mL, Sigma) for 6–48 h. Sodium arsenite (NaAsO₂, 250 µM, 1 h, Merk) served as a positive control for SG formation. In selected experiments, cycloheximide (CHX, 200 µM, Sigma) was used 24 h prior NaAsO₂ and concomitant to LPS treatment, to prevent SG assembly by blocking translation elongation.

### Small RNA Extraction, Library Preparation, and Sequencing

For RNA extraction from cartilage, 150 mg of frozen tissue were pulverised, homogenised in QIAzol (Qiagen) and purified using the miRNeasy Mini Kit (Qiagen). Small RNA libraries were generated using the TruSeq Small RNA Library Preparation Kit (Illumina).

Total RNA was isolated from primary chondrocytes using the miRNeasy Micro Kit (Qiagen)^16^. To improve ligation and reverse transcription efficiency of modified tRNA fragments, RNA was pre-treated using the rtStar™ tRF&tiRNA Pretreatment Kit (Arraystar). Libraries were prepared with the QIAseq miRNA Library Kit^16^.

Library quality was assessed with the Agilent DNA 1000 assay and sequencing was performed on Illumina NovaSeq6000 and HiSeq2000 platforms (SE, 100 cycles).

### Computational Profiling of tRFs

Small RNA seq reads were quality-checked, UMI-collapsed using CLC Genomics Workbench v23.0.5 or adapter-trimmed using Cutadapt v5.1 on Galaxy^17^. tRFs were identified with MINTmap, filtered with Threshold-seq v1.1 and profiled using MINTmap v1.0^18^, retaining only fragments uniquely assigned to the tRNA space. Low-abundance features were removed with Thresholdseq (v1.1). Differential expression analysis was performed using DESeq2 (R package v1.42.1)^19, 20^. tRFs with |FC| ≥ 2 and FDR ≤ 0.1 were considered significantly modulated. The RNA-seq datasets were deposited in GEO (GSE316834; GSE316835). Rank-rank hypergeometric overlap between datasets was assessed with RedRibbon (v1.3.1.)^21^.

### Target Retrieval and Pathway Enrichment

AGO2 CLIP data-supported targets of LPS-modulated tRFs were obtained with tRF-Tar^22^. Gene symbols were mapped to stable Ensembl IDs using GeneCards and ShinyGO^23^. Target sets for up-and down-regulated tRFs were compared using InteractiVenn, retaining only condition-specific targets. Functional enrichment for Gene Ontology (GO) Biological Process (BP), Cellular Component (CC) and Molecular Function (MF) terms was performed using ShinyGO v0.82. Significant pathways were selected using FDR ≤ 0.1.

### Small RNA Expression Analysis by qPCR

Real-Time qPCR was performed using the miRCURY LNA system (Qiagen) with cDNA from 100 ng total RNA and LNA-enhanced primers specific for miRNAs or custom-designed for tRFs using SYBR Green chemistry. Expression levels were normalized to the geometric mean of Ct values from three reference small RNAs (U6 snRNA, RNU48, and let-7-5p), selected for their stability. Relative expression was calculated using the ΔΔC_t_ method.

### Protein Fractionation and Western Blotting

NP40-soluble and NP40-insoluble fractions were prepared following a workflow adapted from^24, 25^. Cells were lysed in cold NP40 lysis buffer, centrifuged and the resulting supernatant was collected as the NP40-soluble fraction. The pellet was resuspended in SDS lysis buffer and processed as the NP40-insoluble fraction. Equal protein amounts were separated by SDS-PAGE and transferred to PVDF membranes. After blocking, membranes were incubated with the following primary antibodies: anti–Pan-Actin (Invitrogen, MA5-11869; 1:2000), anti-Calnexin (Santa Cruz, sc-11397; 1:500), anti–G3BP1 (Santa Cruz, sc-365338; 1:1000), anti–HSP70 (Santa Cruz, sc-515873; 1:1000), anti–Ubiquitin (Santa Cruz, sc-166553; 1:1000), anti–AGO2 (Abcam, ab186733; 1:1000), and anti–HuR (Santa Cruz, sc-5261; 1:1000), HRP-conjugated secondary antibodies (anti-mouse #115–035-174, 1:5000 or anti-rabbit #211-032-171, 1:10,000 Jackson ImmunoResearch).

### Stress Granule Detection by Immunofluorescence

Cells were fixed in 4% PFA (Sigma) and post-fixed in cold methanol. After blocking with 0.1 % (v/v) Triton X-100 in Tris-buffered saline (TBS) and 5% bovine serum albumin for 30 min, samples were incubated with anti-AGO2, anti-G3BP1, or anti-HuR antibodies (1:100) and 15 ug/mL Alexa Fluor 488 goat anti-rabbit (Invitrogen, #A11008) or Cy3 sheep anti-mouse IgG (Sigma, #c2181) conjugated secondary antibodies. Nuclei were stained with DAPI (Merck). Images were acquired using Evident IX83 inverted microscope equipped with the Evident CellSens Dimension software.

### Antisense Oligonucleotide Transfection

C28/I2 chondrocytes were seeded at a density of 2.5 × 10⁵ cells per well in 6-well plates. ASOs targeting 3’tRF^Asp(GTC)^ or scramble controls (Qiagen) were transfected into C28/I2 cells at 25 nM using Lipofectamine™ RNAiMAX (Invitrogen). The cells were treated with LPS the following day and collected 24 hours later.

### mRNA Expression Analysis

Total RNA from ASO-treated C28/I2 cells was reverse-transcribed using the Wonder RT Kit (Euroclone). Real-Time qPCR was performed using TB Green Premix Ex Taq II (Diatech) for genes associated with inflammation and catabolism (*COX2*, *MMP13*). *GAPDH* was used as reference.

### RNA Immunoprecipitation

RNA immunoprecipitation (RIP) was performed to assess the association of small RNAs with AGO2 and to evaluate LPS-induced enhancement of RNA-induced silencing complex (RISC) activity. Cells were lysed in polysome buffer. Lysates were cleared and incubated for 2 h at 4 °C with Protein A/G magnetic beads (Pierce, Thermo Fisher) pre-bound to 5μg AGO2 antibody (Abcam, ab186733) or 5μg IgG control (Thermo Fisher, #026102). After washing, complexes were eluted and RNA was extracted. 10% of each lysate was saved as input for Western blot and 10% for qPCR. RNA enrichment for specific miRNAs and 3’tRF^Asp(GTC)^ was quantified using small-RNA qPCR assays. RIP efficiency and specificity were verified by AGO2 Western blotting.

### Polysome Profiling Coupled to qPCR

To analyze translational regulation upon LPS treatment or tRF modulation, polysome profiling was performed following established procedures^26, 27^. Cell pellets were lysed with polysome buffer, added with 200µM CHX to stabilize ribosome–mRNA complexes and centrifuged. Cytoplasmic extracts were layered onto 10–50% sucrose gradients and ultracentrifuged at 222,300*g* for 3 h. Thirty-four fractions were collected and absorbance at 254 nm was registered. RNA from each fraction was extracted, reverse-transcribed, and subjected to Real-Time qPCR analysis of *MMP13*, *COX2*, and 3’tRF^Asp(GTC)^. Fraction identity and separation quality were verified using Real-Time qPCR for 18S and 28S rRNA.

### *In silico* analysis of 3’tRF^Asp(GTC)^–RBP interactions and functional enrichment

To investigate potential RBP-mediated functions of cartilage-associated tRFs, an exploratory analysis was performed using catRAPID omics v2.1^28^. Based on MINTbase annotation, the mature parental tRNA sequence (tRNA^Asp(GTC)^1-1) of 3’tRF^Asp(GTC)^ was retrieved from gtRNAdb^29^, and used as input. RBPs predicted to bind regions matching the fragment coordinates were ranked by their RNA-binding propensity scores and analysed by pre-ranked GSEA (v4.4.0) across GO BP, CC and MF gene sets. Significant categories (p-value ≤ 0.05) were visualised as scatter dot plots using ggplot2 (R Package v3.5.2)^30^.

### Statistical Analysis

Data were collected from at least three independent experiments. Results are presented as mean values ± standard error of the mean (SEM). Statistical analyses were performed using parametric Student’s t-test for comparisons between two groups. For comparisons among more than two groups, data distribution was assessed using a normality test and followed by one-way ANOVA with Sidak post hoc correction test. Statistical significance was set at FDR ≤ 0.1 or p-value ≤ 0.05. The coefficient of determination (R²) and Spearman correlation for Real-Time qPCR analyses and body mass index (BMI) were calculated. All analyses and graphical representations, including volcano and MA plots, were performed using GraphPad Prism v10.0 (GraphPad Software, San Diego, CA, USA). tRF class distribution was assessed using 2×2 contingency tables and two-sided Fisher’s exact tests on differentially expressed fragments (|FC| ≥ 2, p ≤ 0.05) in LPS-treated chondrocytes and OA cartilage compared to controls.

## Results

### tRF modulation occurs in OA cartilage and is in part mediated by inflammatory stimuli

We analyzed small RNA seq data from OA patient-*versus* healthy donor-derived cartilage (**Fig.1A**). Differential expression analysis identified 672 deregulated tRFs (|FC| ≥ 2 and FDR≤ 0.1), whose 488 significantly upregulated and 184 downregulated (**Fig.1B**). To investigate whether the tRF modulation observed in patients was due at least in part to environmental inflammatory cues characteristic of metabolic OA, we performed small RNA sequencing on primary cultures stimulated with LPS for 6 h and compared each treated sample with its matched untreated control (**Fig.1A**). Interestingly, LPS significantly induced tRNA cleavage with 22 tRFs upregulated (|FC| ≥ 2 and FDR≤ 0.1) (**Fig.1C**). Class-level analysis revealed a selective enrichment of specific tRF classes in both datasets, with 3’tRFs significantly overrepresented in OA cartilage and LPS-treated chondrocytes. 5’tRNA-halves were found to exclusively upregulated in both OA cartilage and LPS treatment (**Fig. 1D–E**; **Fig. S1A-B**; **Fig. S2**). To compare the tRF signatures of OA patient cartilage and the *in vitro* inflammation model, we performed a rank–rank hypergeometric overlap (RRHO) analysis, which revealed a strong overlap in the upper-right quadrant. This region comprises 149 commonly upregulated tRFs in both datasets. Among these, 73 tRFs displayed FC >2, 31 tRFs showed FC >2 and p <0.05, and 3 tRFs passed the multiple testing correction (FDR <0.1) (**Fig. 1F**), naming 5’tRF^Glu(CTC)^, 3’tRF^Asp(GTC)^ and 5’tiRNA^Glu(CTC)^). Collectively, these results indicate that tRF biogenesis mechanisms occurring in OA patient cartilage can be partially recapitulated *in vitro* by LPS exposure of chondrocytes.

**Figure 1.**
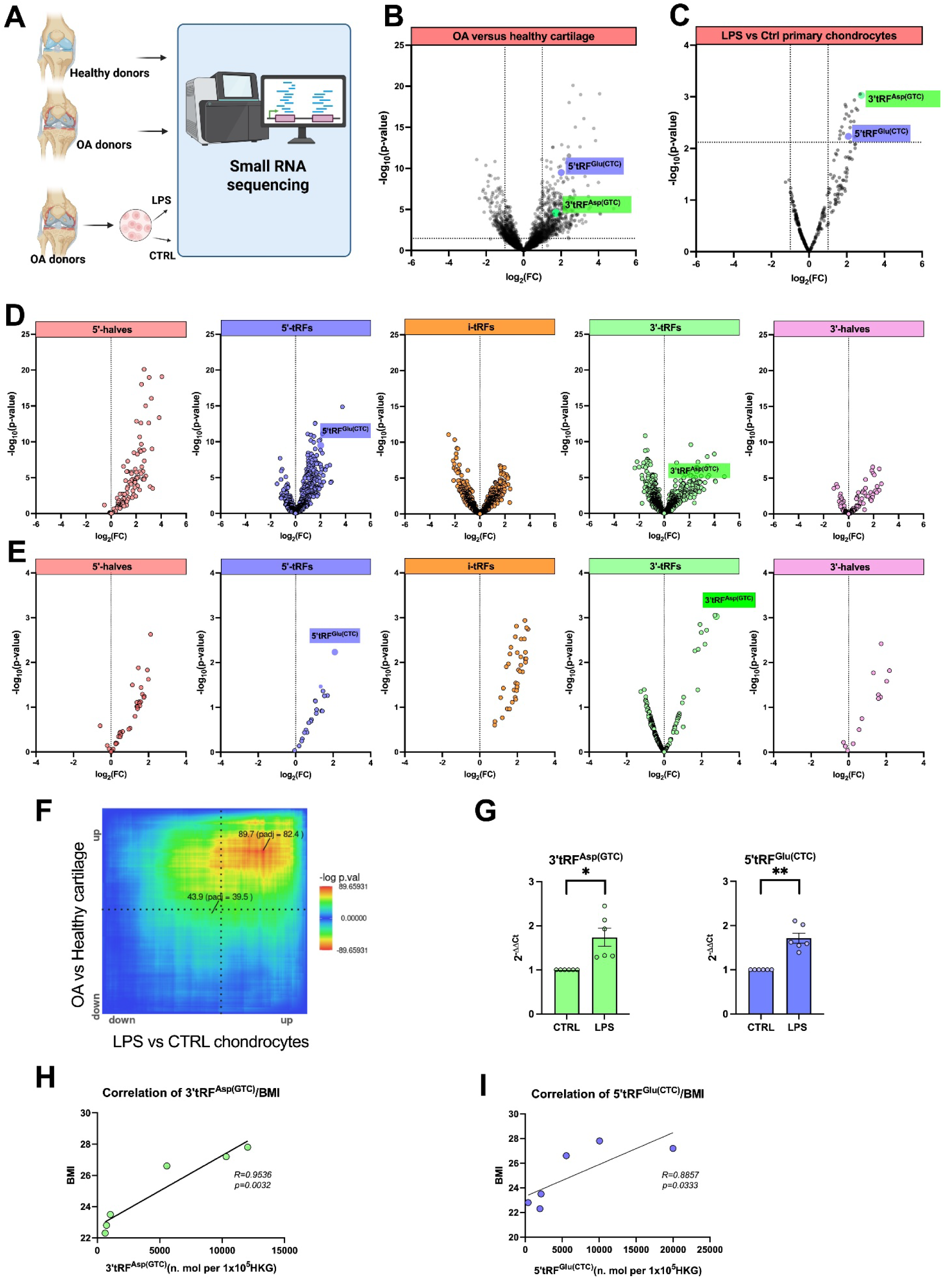
tRF signature in OA cartilage and human primary chondrocytes upon LPS treatment. Small RNA sequencing was performed on cartilage from OA patients and healthy donors and on OA patient–derived primary chondrocytes treated with lipopolysaccharide (LPS, 10 μg/mL, 6 h) (A). Differential expression analysis of DESeq2-normalised tRNA-derived fragment (tRF) counts compared OA (n = 6) versus healthy cartilage (n = 7) (B) and LPS-treated chondrocytes (n = 3) versus matched controls (n = 3) (C). The modulation of specific tRF classes is shown for OA cartilage (D) and LPS-treated chondrocytes (E), highlighting 5’tRF^Glu(CTC)^ and 3’tRF^Asp(GTC)^. Rank-rank hypergeometric overlap analysis between OA cartilage and LPS-treated chondrocytes was performed using RedRibbon (F). Candidate tRFs were validated by Real-Time qPCR in LPS-treated chondrocytes (n = 6) (G) and correlated with body mass index (BMI) using Spearman analysis (H–I). *p < 0.05, **p < 0.01 by Student’s t-test.

To further elucidate the impact of tRF as post-transcriptional regulators, we next explored their potential miRNA-like function. We focused on the LPS-driven tRF signature in order to avoid the pathophysiological complexity and narrow the analysis to inflammatory mechanisms. Applying a less strict statistical threshold of p<0.05, we obtained a dataset of 371 upregulated and 24 downregulated tRFs due to LPS exposure. Only genes predicted to be targeted exclusively by upregulated or downregulated tRFs were considered. 235 genes with potential target sites shared by tRFs modulated in both directions were excluded by the analysis. (**Fig. S3A**). Pathway enrichment analysis revealed distinct regulatory signatures for upregulated and downregulated species. Targets of upregulated tRFs were enriched in intracellular processes, stress responses, and cell death, with functions linked to adhesion, RNA–protein complexes, vesicles, and ribosomal or binding activities. (**Fig.S3B1-3**). In contrast, downregulated tRF targets displayed a more restricted enrichment profile. Identified terms highlighted Golgi reassembly, cell–cell adhesion, mitochondrial membranes, and extracellular vesicle compartments (**Fig.S3C1-3**). Together, these enrichment patterns suggest that LPS drives a coordinated reconfiguration of tRF-mediated miRNA-like regulatory networks in chondrocytes, with upregulated tRFs promoting stress-adaptive and proteostasis-related programs, while downregulated tRFs may relieve constraints on mitochondrial and cell-cell communication pathways.

We then identified potential tRF candidates for downstream functional validation, focusing on their modulation and clinical relevance. In particular, tRFs corresponding to 3’tRF^Asp(GTC)^ and 5’tRF^Glu(CTC)^ were consistently upregulated in OA patient cartilage as well as in LPS-exposed chondrocytes, passing FC and FDR thresholds (highlighted in Fig.1B-E). LPS-mediated induction of 3’tRF^Asp(GTC)^ and 5’tRF^Glu(CTC)^ was validated by Real-Time qPCR in both primary chondrocytes and C28/I2 cells (**Fig.1G**; **Fig.S4A**). In the context of a metabolically driven OA phenotype, body mass index (BMI) represents an established clinical proxy for obesity-associated metabolic stress and metabolic dysfunction linked to osteoarthritis ^1, 2^. On this basis, we investigated whether tRF expression levels are associated with BMI in OA chondrocytes. Spearman correlation analysis revealed a positive association between BMI and 3’tRF^Asp(GTC)^ and 5’tRF^Glu(CTC)^ levels in OA chondrocytes (**Fig.1H-I**), suggesting a potential link between metabolic status and tRF regulation.

### 3’tRFAsp(GTC) inhibition prevents OA-related cellular responses to LPS

Notably, the LPS-treated chondrocyte dataset showed a higher number of distinct tRFs derived from the Asp isoacceptor (**Fig.S4B**). This observation prompted us to focus on 3’tRF^Asp(GTC)^ to investigate its role in mediating LPS responses in OA chondrocytes We used ASO approach to target and inhibit this specific tRF (α3’tRF^Asp(GTC)^), compared with a negative control ASO (αNC). Upon transfection, cells were treated with LPS for 24h (**Fig.2A**). The efficiency of 3′tRF^Asp(GTC)^ silencing was evaluated by qPCR, which confirmed a significant reduction of the fragment level in LPS-treated α3’tRF^Asp(GTC)^ cells (**Fig.2B**). We then measured the gene expression of two OA markers, *COX2* and *MMP13*, already reported to be upregulated by LPS^5^. COX2 is responsible for producing pro-inflammatory prostaglandins, while MMP13 is a matrix metalloproteinase involved in cartilage collagen breakdown during OA. For both *COX2* and *MMP13*, as expected, a strong and significant induction was observed upon LPS stimulation. Interestingly, this upregulation was strongly inhibited in α3’tRF^Asp(GTC)^ cells (**Fig.2C-D**).

**Figure 2.**
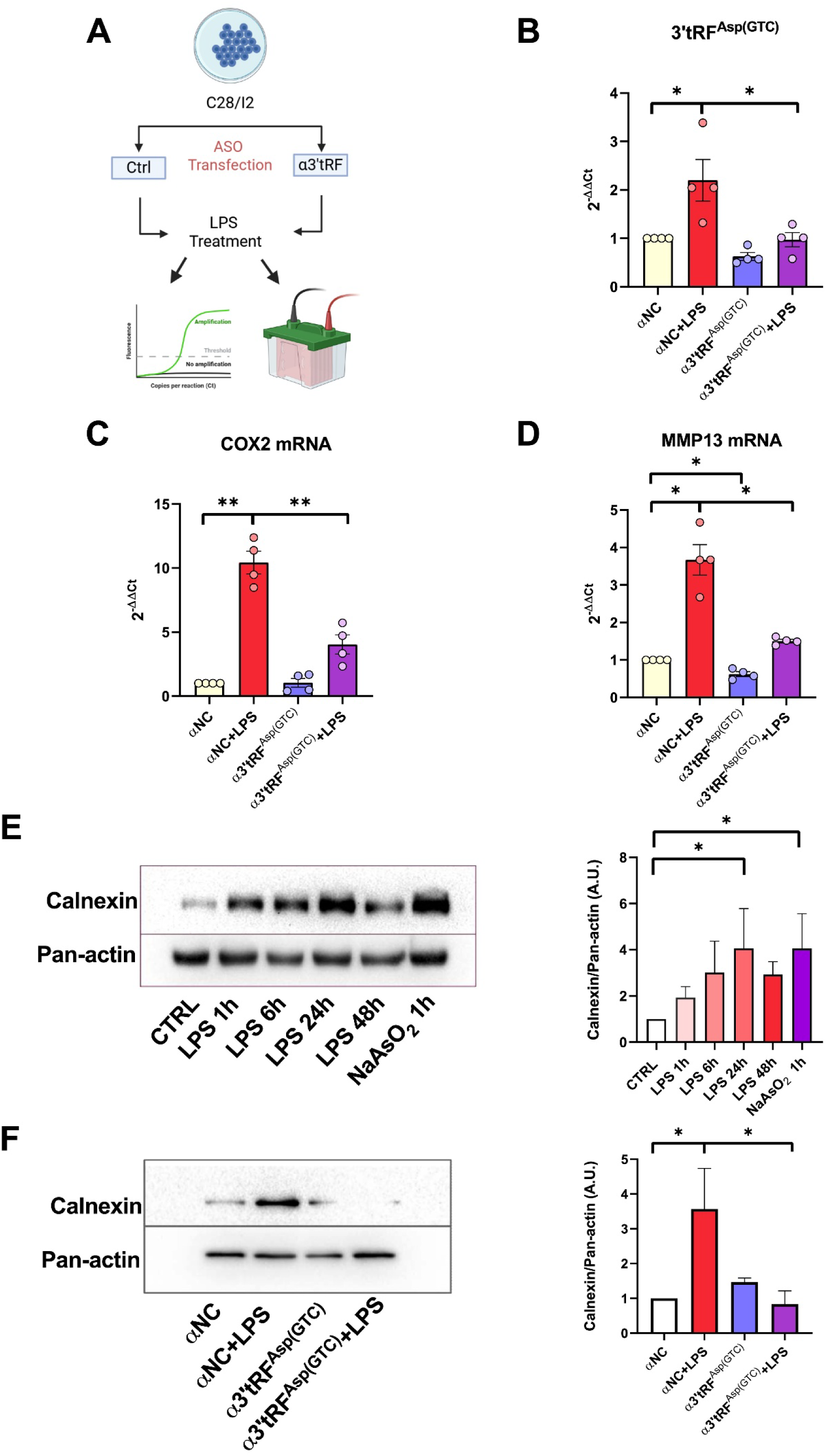
3’tRF^Asp(GTC)^ mediates LPS-induced OA changes. C28/I2 chondrocytes were transfected with an antisense oligonucleotide (ASO) against 3’tRF^Asp(GTC)^ (α3′tRF^Asp(GTC)^) or a scramble negative control (αNC), and subsequently stimulated with LPS for 24 h. Cells were then analysed by Real-Time qPCR and westernblot (A). 3’tRF^Asp(GTC)^ levels were detected and ASO transfection efficiency was confirmed by Real-Time qPCR (B). *COX2* and *MMP13* gene expression were also detected by Real-Time qPCR (C-D). Protein levels of calnexin were measured by westernblot in C28/I2 chondrocytes treated with LPS in time-course. Representative image of westernblot (left) and densitometry quantification were shown (n=3, right) (E). An antisense oligonucleotide (ASO) against 3’tRF^Asp(GTC)^ (α3’tRF^Asp(GTC)^) prevented calnexin upregulation. Representative image of westernblot (left) and densitometry quantification were shown (n=4, right) (F) *p<0.05, **p<0.01 by One-way ANOVA with Sidak correction for multiple comparisons.

Previously, we and others have demonstrated the involvement of unfolded protein response (UPR) systems and ER stress in inflammatory pathologies, including OA^14, 31^. As shown in **Fig.2E**, we confirmed that LPS treatment in C28/I2 chondrocytes upregulates the expression of Calnexin, a crucial ER chaperone whose upregulation correlates with ER stress. The inhibition of 3’tRF^Asp(GTC)^ prevented LPS-induced Calnexin upregulation, suggesting the attenuation of ER stress (**Fig.2F**). Altogether, these results suggest that 3’tRF^Asp(GTC)^ could mediate inflammation-induced catabolism and ER stress pathways in OA.

### 3’tRFAsp(GTC) binds to AGO2 and modulates the assembly of LPS-induced AGO2-positive stress granules

One of the mechanisms by which tRFs mediate gene regulation is the interaction with AGO2 in the RISC complex^32^. Our *in silico* prediction suggests a possible relevant role of tRFs in AGO2–mediated gene regulation. To explore this hypothesis, we performed AGO2-RIP assay. We immunoprecipitated AGO2 from LPS-treated C28/I2 cell extracts and purified the associated RNAs (**Fig.3A**). We found that 3’tRF^Asp(GTC)^ is co-immunoprecipitated with AGO2, and its levels in the AGO2 IP fraction increased in chondrocytes treated with LPS (**Fig.3B**). To validate our AGO2-RIP assay, we assessed miR-155 levels associated with AGO2. This is a known pro-inflammatory miRNA marker in chondrocytes^7^, and a canonical RNA guide within the RISC complex. The miR155-5p was significantly co-immunoprecipitated with AGO2 and its levels increased in chondrocytes treated with LPS (**Fig.3C**). At the same time, the small nuclear RNA U6 was not detected in AGO2 pull-down fraction (**Fig.3D**), corroborating the specificity of the assay.

**Figure 3.**
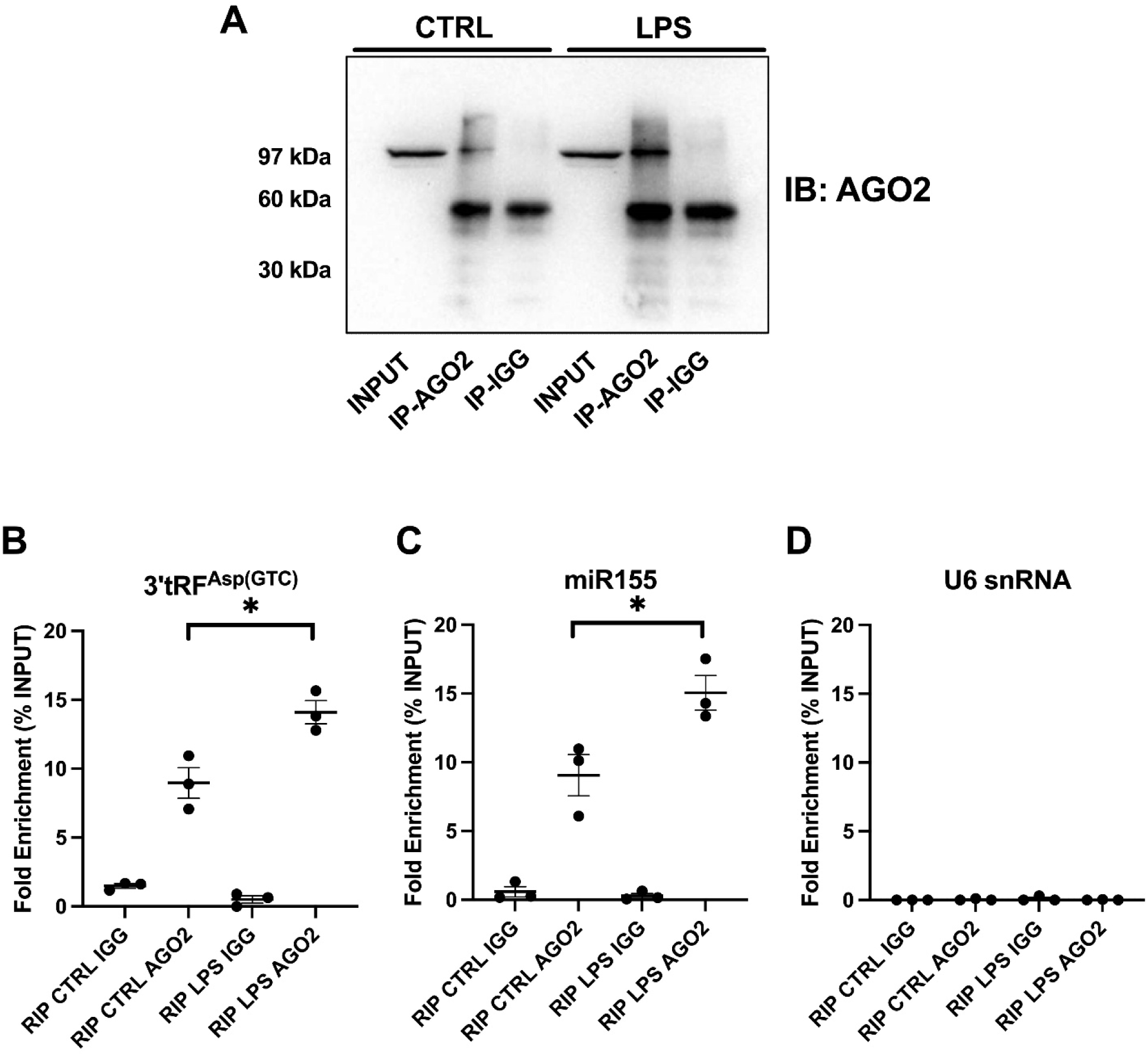
3’tRF^Asp(GTC)^ interacts with AGO2. C28/I2 chondrocytes were treated with 10μg/mL LPS for 24h and AGO2-associated RNAs were processed using an AGO2-RIP assay. Representative western blot showing AGO2 protein immunoprecipitation using an anti-AGO2 antibody. 10% of the input lysate and 10% of the AGO2-RIP eluate were loaded for comparison (A). Real-Time qPCR analysis of 3’tRF^Asp(GTC)^, miR-155 as positive control, and U6 snRNA as negative control was performed in AGO2 immunoprecipitates, expressed as percentage of input, with rabbit IgG used as negative control (B-C). *p<0.05 by One-way ANOVA with Sidak correction for multiple comparisons.

Although the role of tRFs as miRNA-like gene regulators has been previously suggested based on their association with AGO2^33^, the difference in sequence length and structure compared to canonical miRNAs may suggest additional tRF-mediated mechanisms. Since AGO2 localization and function are dynamically regulated under cellular stress conditions, and SGs represent key sites of translational repression, we investigated whether LPS-dependent modulation of 3’tRF^Asp(GTC)^ affects AGO2 localization and possibly SG assembly. NaAsO₂ treatment is known to induce SG formation and thus used in this study as a positive control treatment^15^. We found that LPS exposure induced the redistribution of AGO2 in cytoplasmic foci in C28/I2 chondrocytes. Interestingly these foci were identified as SGs, as they were positive for the SG-specific marker G3BP1^15^. NaAsO₂ treatment induced clear G3BP1-positive foci without colocalization with AGO2. Interestingly, the inhibition of 3’tRF^Asp(GTC)^ prevented the formation of AGO2– and G3BP1-positive foci but only in LPS conditions, suggesting a potential novel mechanism by which this tRF modulates AGO2 function in SG assembly as an inflammatory response (**Fig.4A**; **Fig.S5**). As an independent approach, we performed differential extraction from soluble and insoluble protein fractions followed by Western blot analysis of AGO2 and SG markers (HSP70 and HuR, respectively, as chaperone and RBP consistently recruited into SGs). Also, this approach confirmed that the 3’tRF^Asp(GTC)^ inhibition prevents the inclusion of AGO2 and SG markers in the insoluble fraction (**Fig.4B-C**).

**Fig. 4.**
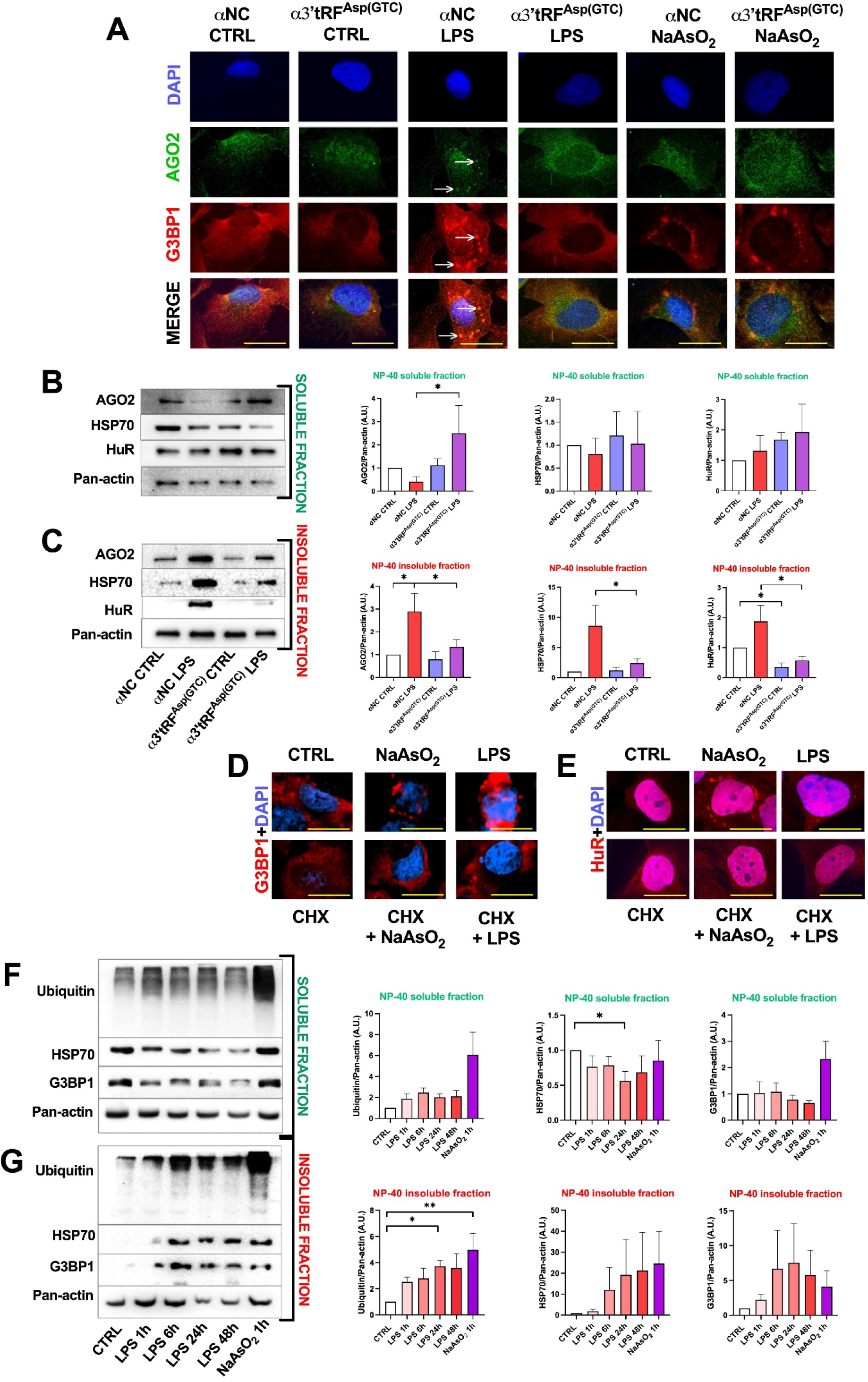
3’tRFAsp(GTC) inhibition prevents AGO2 inclusion in stress granules upon LPS treatment. C28/I2 chondrocytes were transfected with an antisense oligonucleotide targeting 3’tRF^Asp(GTC)^ (α3′tRF^Asp(GTC)^) or a scrambled control (αNC), and stimulated with LPS (10 μg/mL, 24 h) or sodium arsenite (NaAsO₂, 250 μM, 1 h). Immunofluorescence analysis was performed to assess co-localization of G3BP1 (red) and AGO2 (green) (A). White arrows indicate G3BP1-AGO2 positive cytoplasmic *foci*. Nuclei were counterstained with DAPI (blue). Representative images were acquired at 60× magnification, digitally enlarged for clarity; scale bar: 20 μm. Representative western blots (left) and densitometric analyses (right) of soluble (B) and insoluble (C) fractions were performed to detect AGO2, HSP70 and HuR, with Pan-actin as loading control (n=4). C28/I2 chondrocytes were treated with LPS or NaAsO₂ as SG-positive control, and/or cycloheximide (CHX, 200 μM) as SG inhibitor. Immunofluorescence analysis of G3BP1 (D) and HuR (E) localization was performed. Representative western blots and densitometric analyses of soluble (F) and insoluble (G) fractions were used to assess polyubiquitinated proteins, HSP70 and G3BP1 following LPS time-course or NaAsO₂ treatment (n=4). *p < 0.05, **p < 0.01 by one-way ANOVA with Sidak correction.

Next, we assessed the nature of observed cytoplasmic foci as SGs by co-treating LPS and NaAsO₂ with CHX, as an inhibitor of translational elongation preventing ribosome run-off and retaining mRNAs within polysomes. Both SG markers, G3BP1 and HuR^15^, redistributed into cytoplasmic foci, whereas co-treatment prevented foci formation, confirming their *bona fide* SG nature (**Fig. 4D-E**; **Fig.S6**).

Moreover, we performed protein differential extraction followed by Western blot analysis of G3BP1, HSP70 and ubiquitinated proteins at multiple time points (1–48 h) of LPS and after NaAsO₂ (**Fig. 4F–G**). G3BP1 progressively shifted from the soluble to the insoluble fraction during LPS stimulation, in line with its accumulation in SGs. HSP70 showed reduced solubility after prolonged stress, consistent with its recruitment to condensates involved in SG remodelling. Poly-ubiquitinated proteins increased particularly in the insoluble fraction after LPS or NaAsO₂ exposure, indicating proteotoxic stress and reduced protein clearance. Interestingly, the accumulation of ubiquitin in both the soluble and the insoluble fractions suggests a potential role for SGs beyond their canonical regulatory functions on RNA, acting as transient hubs where unfolded or aggregation-prone proteins may be sequestered before refolding or degradation. Overall, these data support a stress-dependent redistribution of gene regulatory factors into dynamic, insoluble condensates.

### Inflammatory stress affects translation efficiency by reprogramming chondrocyte translatome

tRFs have been previously described to modulate protein translation in yeast and archaeal organisms^34, 35^, however the molecular basis in higher eukaryotes of this observation is lacking. To further investigate the impact of LPS-induced SG assembly on translational efficiency, we performed polysome profiling coupled with Real-Time qPCR in C28/I2 chondrocytes treated with LPS or NaAsO_2_ (**Fig.5A**). Our results revealed that LPS induces a massive translational block, characterized by polysome collapse, yet displaying a profile distinct from the positive control NaAsO_2_ and puromycin, both known to block ribosomes in monosome fractions^36^ (**Fig.5B1-3**; **Fig.S7**). Fraction identity and separation quality were verified using Real-Time qPCR for 18S and 28S rRNA (**Fig.5C**). Analysis of the area under the curve (AUC) showed a clear increase in free RNA fractions and consistent reduction of heavy polysomes and thus active translation, as well as of light polysomes following LPS treatment (**Fig.5D**). Subsequent Real-Time qPCR on transcripts of the different fractions indicated that the housekeeping gene *GAPDH* exhibited reduced association with polysomes, reflecting a global decreased translation, whereas inflammatory and catabolic genes such as *COX2* and *MMP13* were enriched in polysomal fractions under LPS exposure (**Fig.5E-F**).

**Fig. 5.**
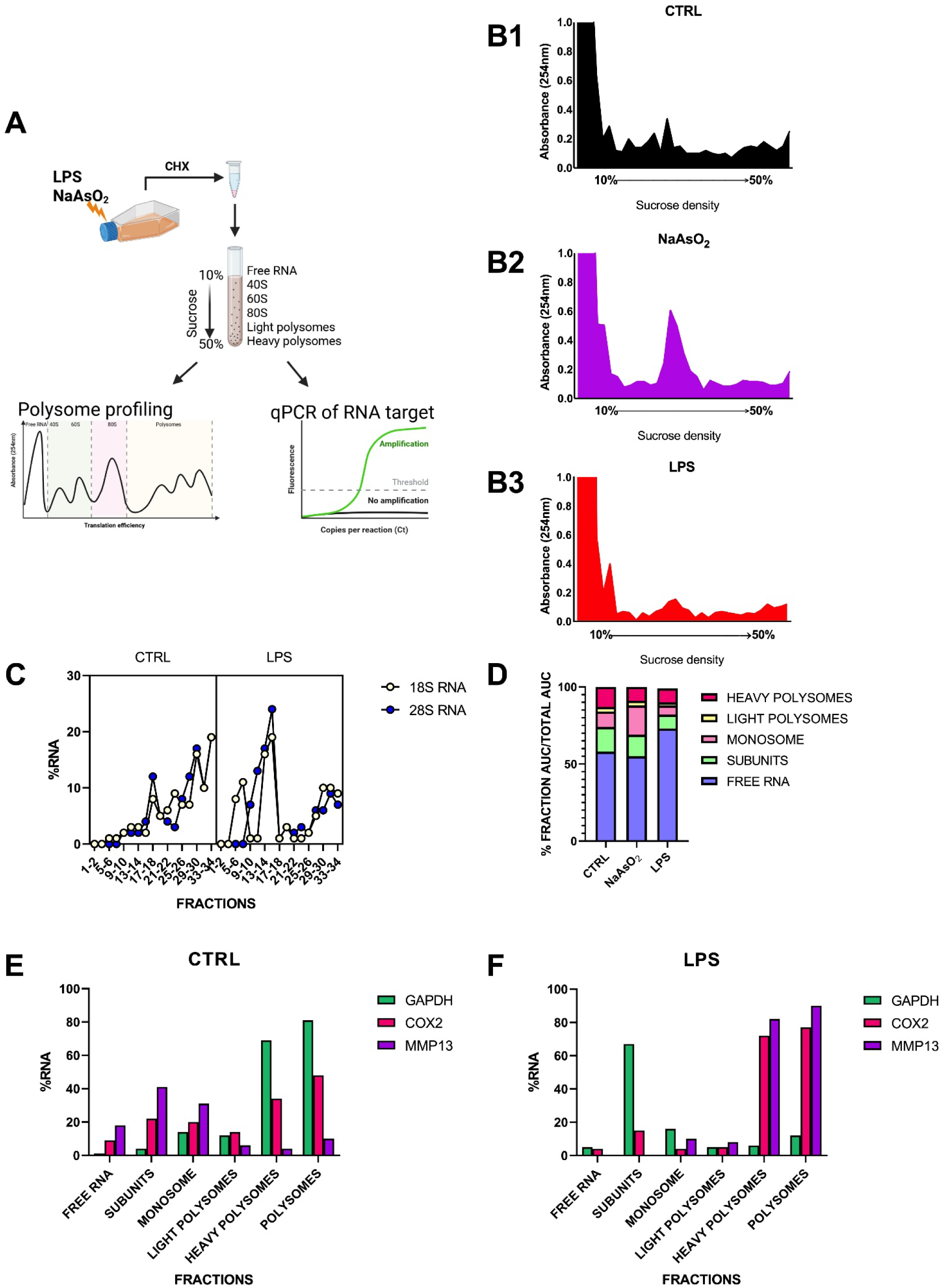
Inflammatory stress affects translation efficiency by reprogramming chondrocyte translatome. Polysome profiling combined with Real-Time qPCR in C28/I2 chondrocytes treated with LPS (10 µg/mL, 24 h) or NaAsO₂ (250 µM, 1 h). Identical lysates were loaded onto 10%–50% sucrose gradients, and polysomes were separated by ultracentrifugation. Each fraction was profiled for RNA content and Real-Time qPCR (A). Sucrose-gradient absorbance profiles are shown for all conditions (B1-3). Fraction identity and separation quality were assessed by Real-Time qPCR for 18S and 28S rRNA (C). Area-under-the-curve (AUC.) analysis was performed for selected polysome fractions and expressed as percentage of the total (D). 40S and 60S indicate ribosomal subunits, 80S monosomes; light polysomes reflect low ribosome occupancy, whereas heavy polysomes indicate active translation. RNA from successive paired fractions was pooled, extracted, and *GAPDH* (housekeeping), *COX2* (inflammatory), and *MMP13* (catabolic) transcripts were quantified by Real-Time qPCR in CTRL and LPS conditions (E–F). Polysome profiles, AUC analyses, and Real-Time qPCR measurements are representative of experiments conducted in triplicate biological replicates.

### 3’tRFAsp(GTC) might mediate LPS-induced blockade of translation initiation by interfering with 40S subunit

To further dissect the role of 3’tRF^Asp(GTC)^ in directly driving LPS-mediated translational modulation, we investigated *in silico* if 3’tRF^Asp(GTC)^ could interact with RBPs involved in the protein quality control system. We employed catRAPID omics 2.1 to predict protein interactors of the fragment and a pre-ranked GSEA was performed (**Fig.6A**). The GO BP enrichment included translation-related processes, mRNA metabolism, stress and inflammatory responses, synaptic regulation. The GO CC enrichment revealed RNP complexes covering multiple steps of RNA metabolism. Terms related to mitochondrial structures and cell–substrate junction were also listed as enriched. The GO MF enrichment included RNA binding and ribosomal functions, ubiquitin-related activities and enzymatic transfer functions (**Fig.6B**). The enrichment of ribosomal components, translation-related functions, and RNA-binding activities suggests that the fragment may interfere with protein synthesis at both initiation and elongation levels. Interestingly, Real-Time qPCR on RNA isolated from polysome fractions confirmed that the fragment was enriched in fractions corresponding to the small ribosomal subunit, consistent with the distribution of 18S rRNA and this enrichment was higher in samples derived from LPS-treated cells (**Fig.6C**).

**Fig. 6.**
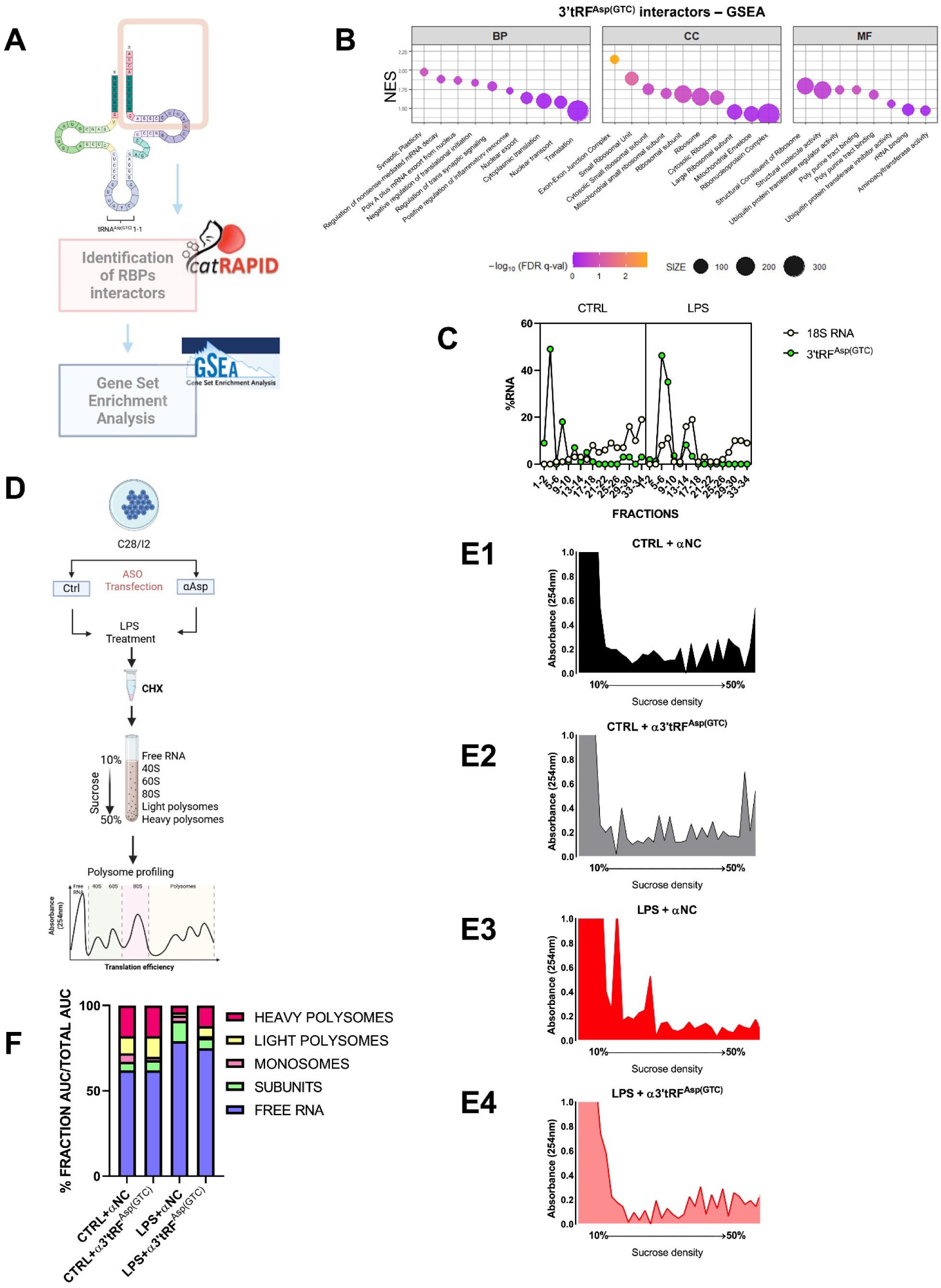
3’tRFAsp(GTC) could mediate LPS-induced blockade of translation initiation by potentially interfering with 40S subunit. Schematic overview of the catRAPID omics 2.1 workflow used to predict protein interactors of 3’tRF^Asp(GTC)^ based on alignment with the parental tRNA and region-restricted interaction scoring, followed by pre-ranked gene set enrichment analysis (GSEA) of predicted interactors (A). Enriched terms within GO Biological Process, Cellular Component and Molecular Function categories are shown (B). RNA from paired sucrose-gradient fractions was pooled, extracted and 3’tRF^Asp(GTC)^ levels were quantified by Real Time qPCR in CTRL and LPS conditions and compared with 18S distribution (C). C28/I2 chondrocytes were transfected with an antisense oligonucleotide targeting 3’tRF^Asp(GTC)^ (α3’tRF^Asp(GTC)^) or a scrambled control (αNC), followed by LPS stimulation for 24 h. Identical lysates were separated on 10–50% sucrose gradients by ultracentrifugation, and fractions were profiled for RNA content (D). Representative sucrose-gradient absorbance profiles are shown (E1–4). Area-under-the-curve (AUC) analysis of selected polysome fractions is expressed as percentage of the total (F). 40S and 60S indicate ribosomal subunits, 80S monosomes; light polysomes reflect low ribosome occupancy, whereas heavy polysomes indicate active translation. Data are representative of three independent biological replicates.

Then, to assess the specific impact of 3′tRF^Asp(GTC)^, we employed ASO strategy in C28/I2 cells in the presence or absence of LPS treatment coupled to polysome profiling (**Fig.6D**). As shown in the profiles of **Fig.E1-4** and the AUC percentage of **Fig.6F**, the inhibition of 3’tRF^Asp(GTC)^ rescues the polysome collapse induced by LPS, suggesting that this fragment is crucial to mediate inflammation-induced translation blockade. These findings indicate that 3’tRF^Asp(GTC)^ potentially associates with small ribosomal subunits and may play a functional role in selective translatome reprogramming.

## Discussion

This study uncovers a broad reprogramming of tRF landscape in OA cartilage. The extensive deregulation observed across multiple tRF classes, with a selective enrichment of 3′– and 5′-derived fragments, supports the existence of regulated tRF biogenesis rather than passive tRNA breakdown. Importantly, a significant fraction of this signature is recapitulated in LPS-stimulated chondrocytes, with a strong overlap between OA cartilage and the *in vitro* inflammatory model. This convergence indicates that inflammatory cues are a major driver of tRF modulation in OA and validates LPS exposure as a relevant system to capture core aspects of inflammation-associated tRF regulation. Within this inflammation-driven tRF program, our findings identify 3’tRF^Asp(GTC)^ as a regulator linking inflammatory stress to translational control and RNP remodelling in chondrocytes. Our findings identify 3’tRF^Asp(GTC)^ as a central regulator linking inflammatory stress to translational control and RNP remodelling in chondrocytes. Rather than a passive degradation product, this tRF functions as an active hub coordinating SG dynamics, ribosome engagement, and AGO2-mediated post-transcriptional repression. These results align with previous studies demonstrating that tRFs regulate post-transcriptional gene expression in chondrocytes: tRF-3003a (3’tRF^Cys(GCA)^) modulates JAK3 via AGO/RISC under IL-1β stimulation, and tRF-5009A (5’tRF^Val(CAC)^) controls autophagy and cartilage degeneration by targeting mTOR, illustrating tRFs as modulators of inflammatory and metabolic pathways in OA^33, 37^. Moreover, SGs have been shown to sequester *COX-2* mRNAs, delaying translation in IL-1β-stimulated chondrocytes, emphasizing the role in post-transcriptional control^8^. A recent study also suggests a potential clinical relevance of tRFs in OA, reporting that serum levels of tRF-5022B correlate with disease severity and distinguish OA patients from healthy controls and RA patients^38^.

Beyond human systems, ribosome-associated tRFs (ranc-RNAs) in yeast and stress-induced tRFs in archaea directly interact with ribosomal subunits and aminoacyl-tRNA synthetases, modulating translation efficiency and protein synthesis rates under stress conditions^34, 35^. These observations support a conserved role for tRFs as functional regulators of translation rather than a passive tRNA byproduct. Consistently, 3’tRF^Asp(GTC)^ accumulates in correspondence with the 40S subunit, selectively engages AGO2 under inflammatory conditions, contributing to SG assembly and linking translational restraint with RNP remodelling.

Collectively, these data reveal 3’tRF^Asp(GTC)^ as a critical mediator of stress adaptation in OA cartilage, integrating translational control, AGO2 engagement, and proteostasis systems. Unravelling these post-transcriptional layers not only provides mechanistic insight but also identifies therapeutic opportunities. With the majority of human proteins remaining “undruggable” translation-targeted strategies—including RNA interference, mRNA-based therapeutics, and tRF modulation—represent promising avenues. By uncovering a regulatory axis connecting altered metabolism, RNA biology, and translational fate, 3’tRF^Asp(GTC)^ emerges as a potential RNA target capable of restoring proteostatic balance and attenuating chronic inflammatory stress in degenerative joint disease.

## Limitations

Our findings indicate that tRFs are dynamically shaped by inflammatory cues with focus on 3’tRF^Asp(GTC)^. Although this fragment emerged as a potent regulator of translational and RNP remodelling pathways, other inflammation-responsive tRFs were also modulated. Their roles-besides ribosomal engagement, AGO2 loading, and stress-adaptive translation-remain to be elucidated.

Moreover, our *in vitro* model captures a subset of the metabolic and inflammatory stresses present *in vivo*. Future studies should define the broader tRF landscape across stimuli, mapping structure–function determinants and validating these axes in animal models.

## Acknowledgments

The drawings in this article were created with BioRender.com.

## Supplementary figures

**Fig. S1.**
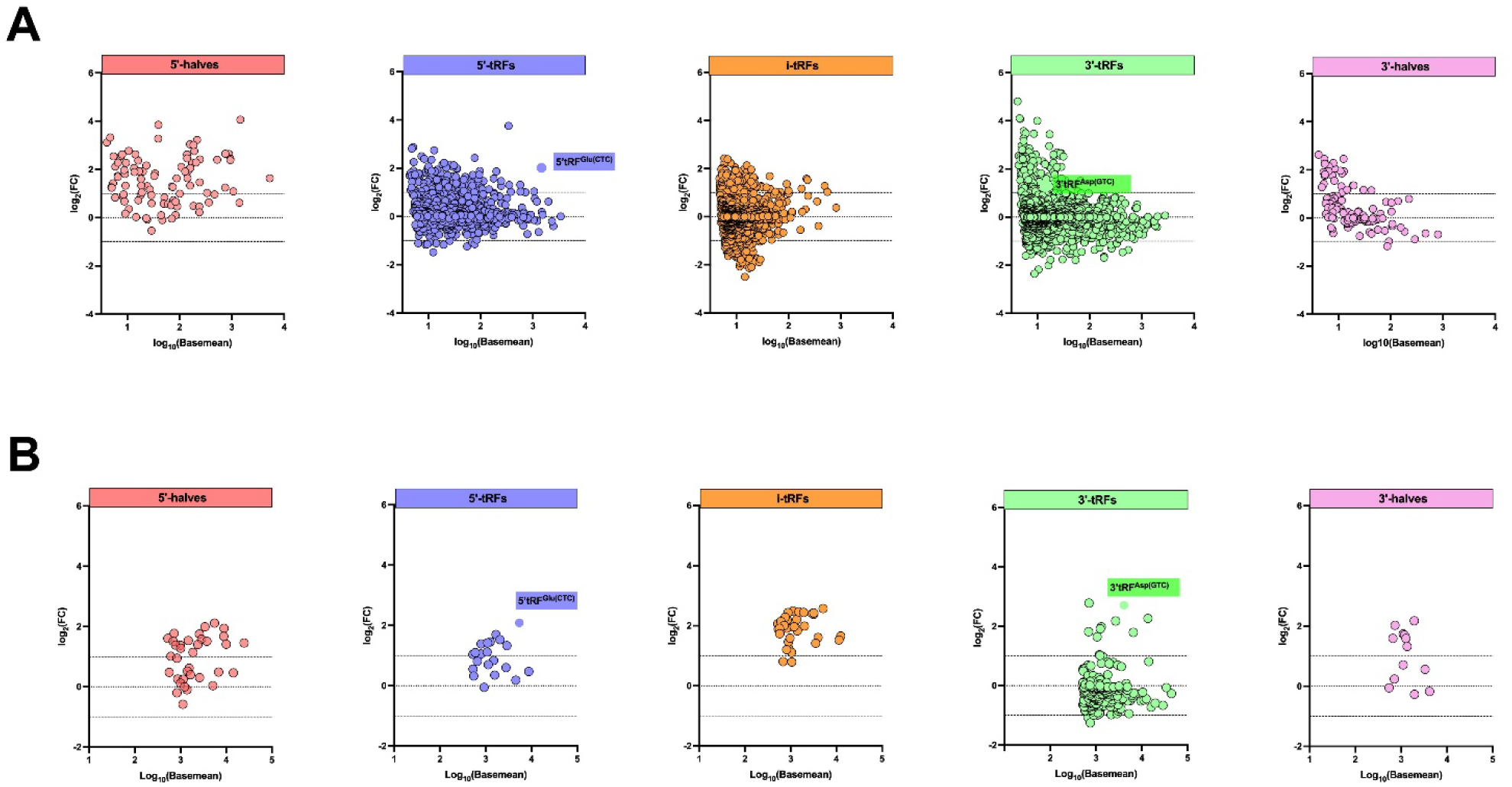
MA plots showed log₂FC as a function of average expression (basemean), highlighting that several abundant fragments in OA cartilage. (A) and OA chondrocyte datasets (B).

**Fig. S2.**
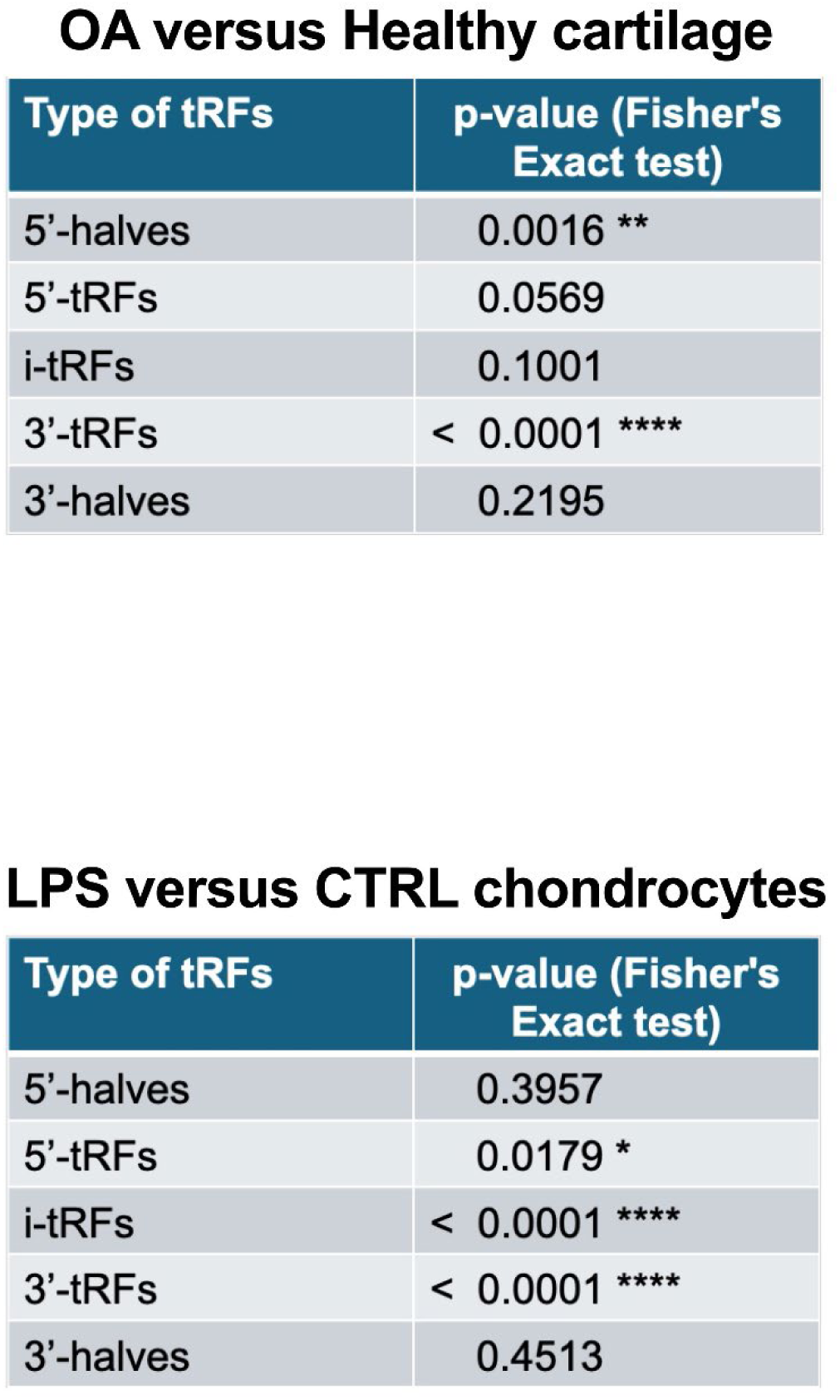
For each tRF class (5′-halves, 5′-tRFs, i-tRFs, 3′-tRFs, and 3′-halves), a 2×2 contingency table was constructed to evaluate whether that class was overrepresented among differentially expressed tRFs (|FC| ≥ 2 and p ≤ 0.05). The number of significant fragments belonging to each class was compared with the number of significant fragments from all other classes, and the same comparison was performed for non-significant fragments. Two-sided Fisher’s exact tests were applied to assess enrichment or depletion of each tRF class in OA cartilage compared to healthy tissue (top panel) and in LPS-treated chondrocytes (bottom panel). Reported p-values correspond to the Fisher test for each class.

**Fig. S3.**
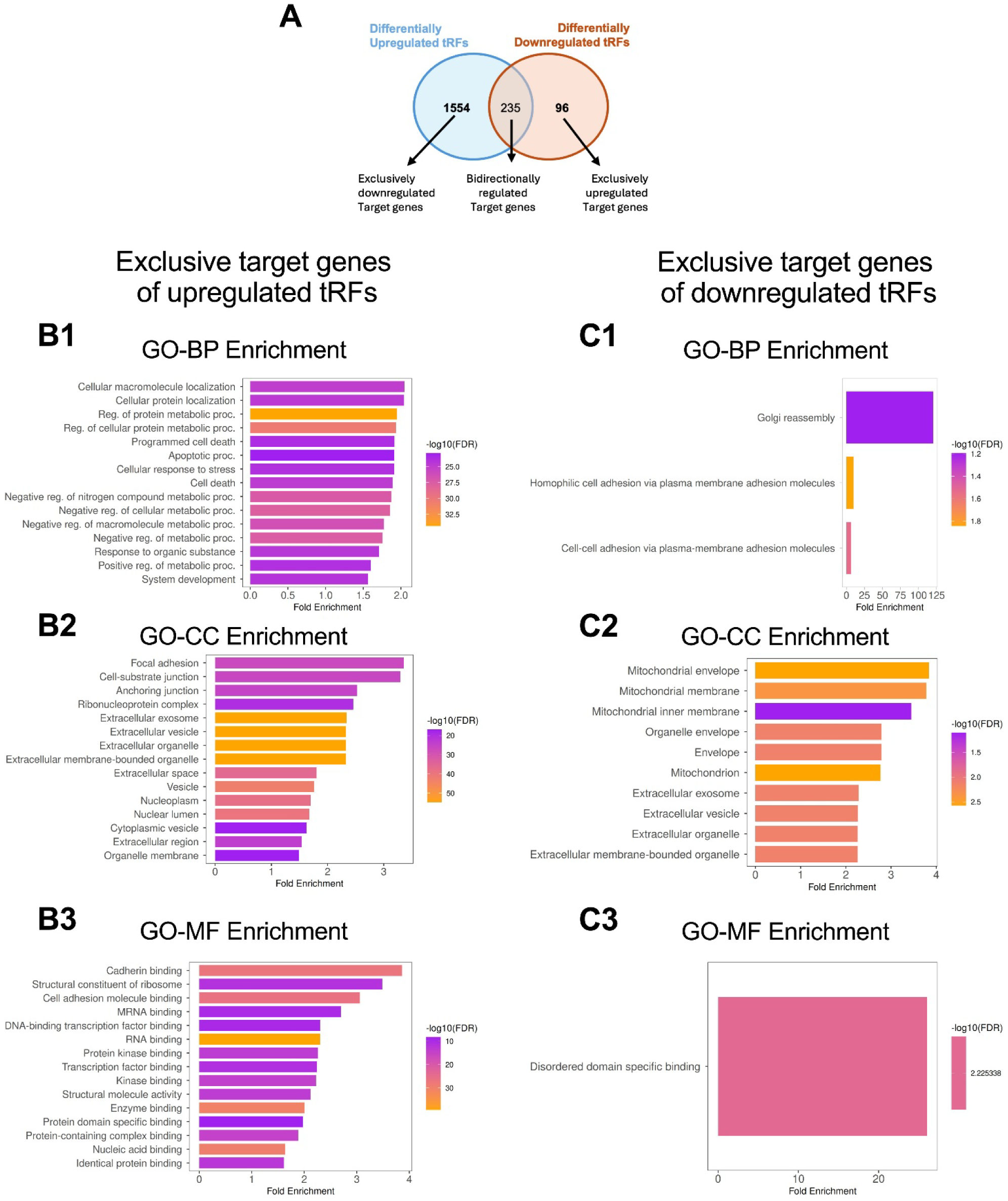
Venn diagram showing the overlap between predicted targets of upregulated (red) and downregulated (blue) tRFs (A). Functional enrichment analyses of exclusive targets of upregulated tRFs: GO Biological Process (B1); GO Cellular Component (B2); GO Molecular Function (B3). Functional enrichment analyses of exclusive targets of downregulated tRFs: GO Biological Process (C1); GO Cellular Component (C2); GO Molecular Function (C3).

**Fig. S4.**
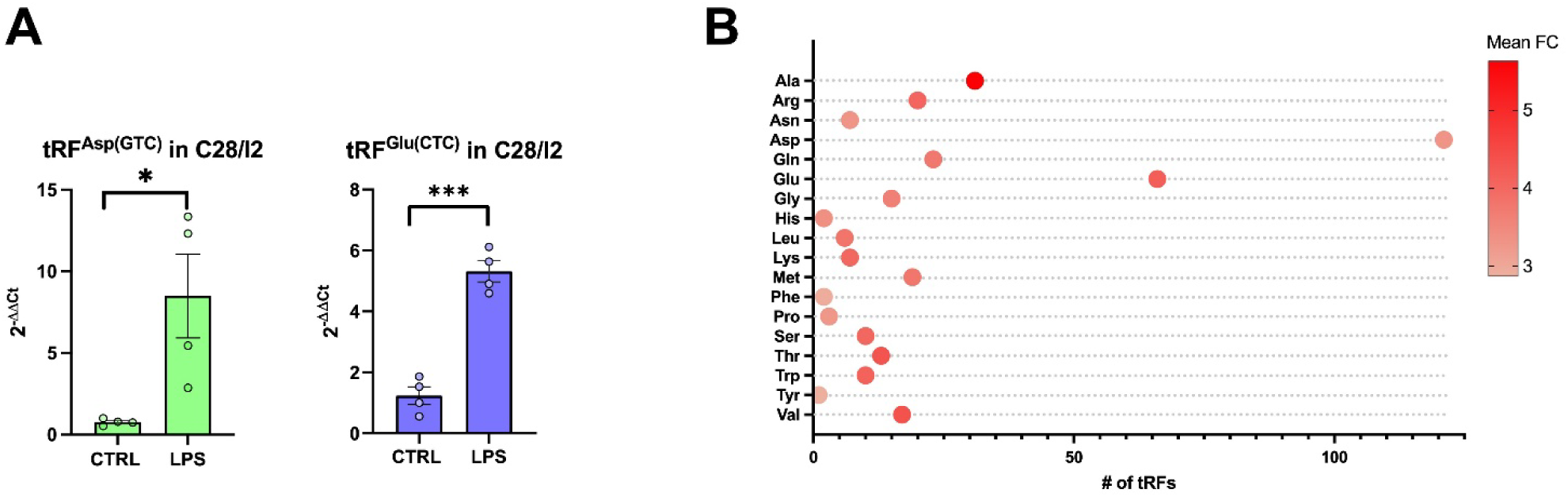
C28/I2 chondrocytes were treated with 10μg/mL LPS for 6 hours and samples collected for RNA isolation. 3’tRF^Asp(GTC)^ and 5’tRF^Glu(CTC)^ levels were measured by Real Time qPCR and comparison analyzed between LPS *versus* untreated control (A). Isoacceptor profiling of tRFs with dot color representing the mean FC (B). *p≤0.05, ***p≤0.001 by Student’s t-test.

**Fig. S5.**
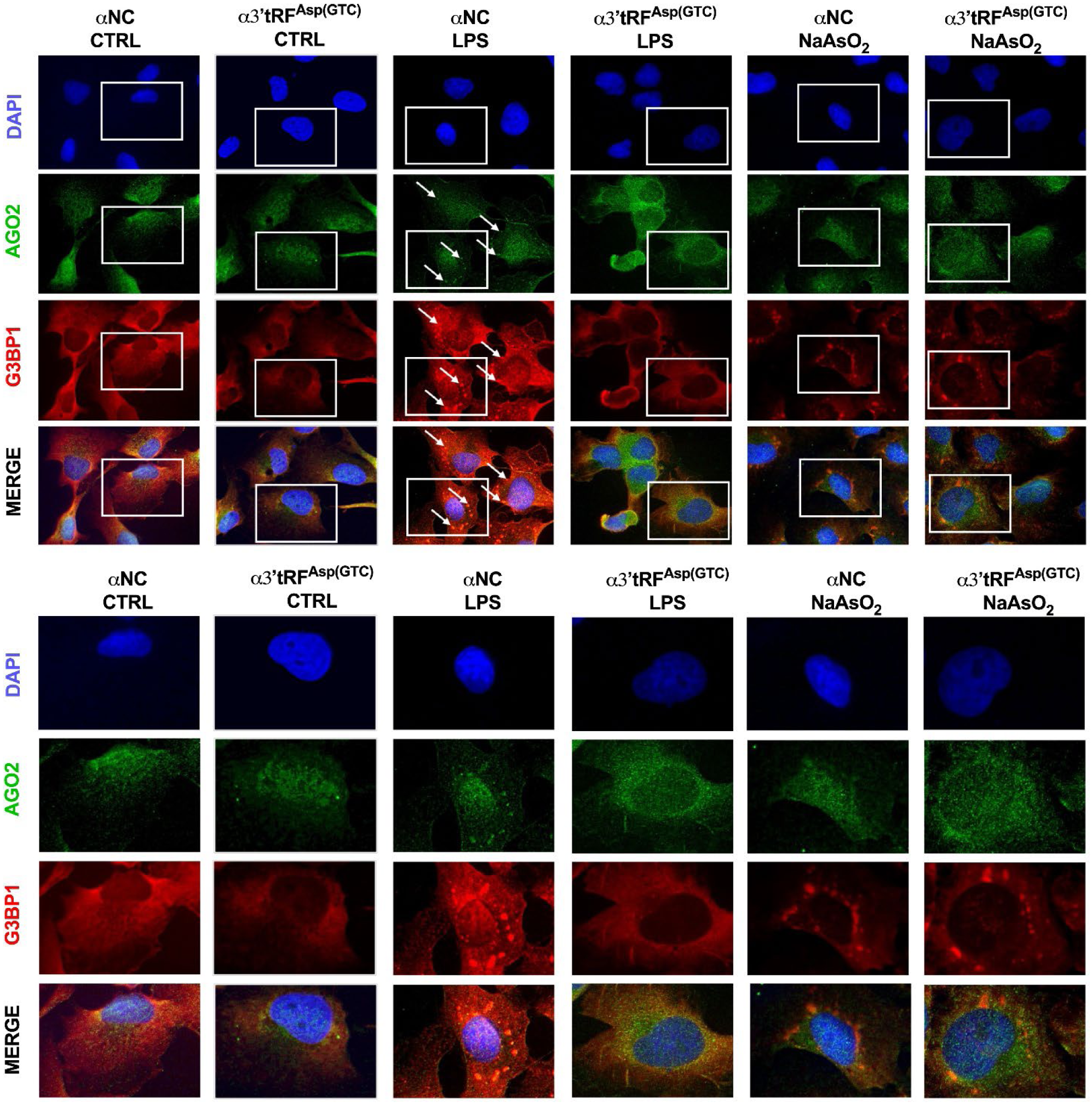
Original immunofluorescence images corresponding to Figure 4A. Representative full-field images showing G3BP1 (red) and AGO2 (green) localization in chondrocytes under the indicated conditions. White boxes indicate the regions selected for magnification and displayed in the main Figure 3A. Nuclei were counterstained with DAPI (blue). Images were acquired at 60× magnification.

**Fig. S6.**
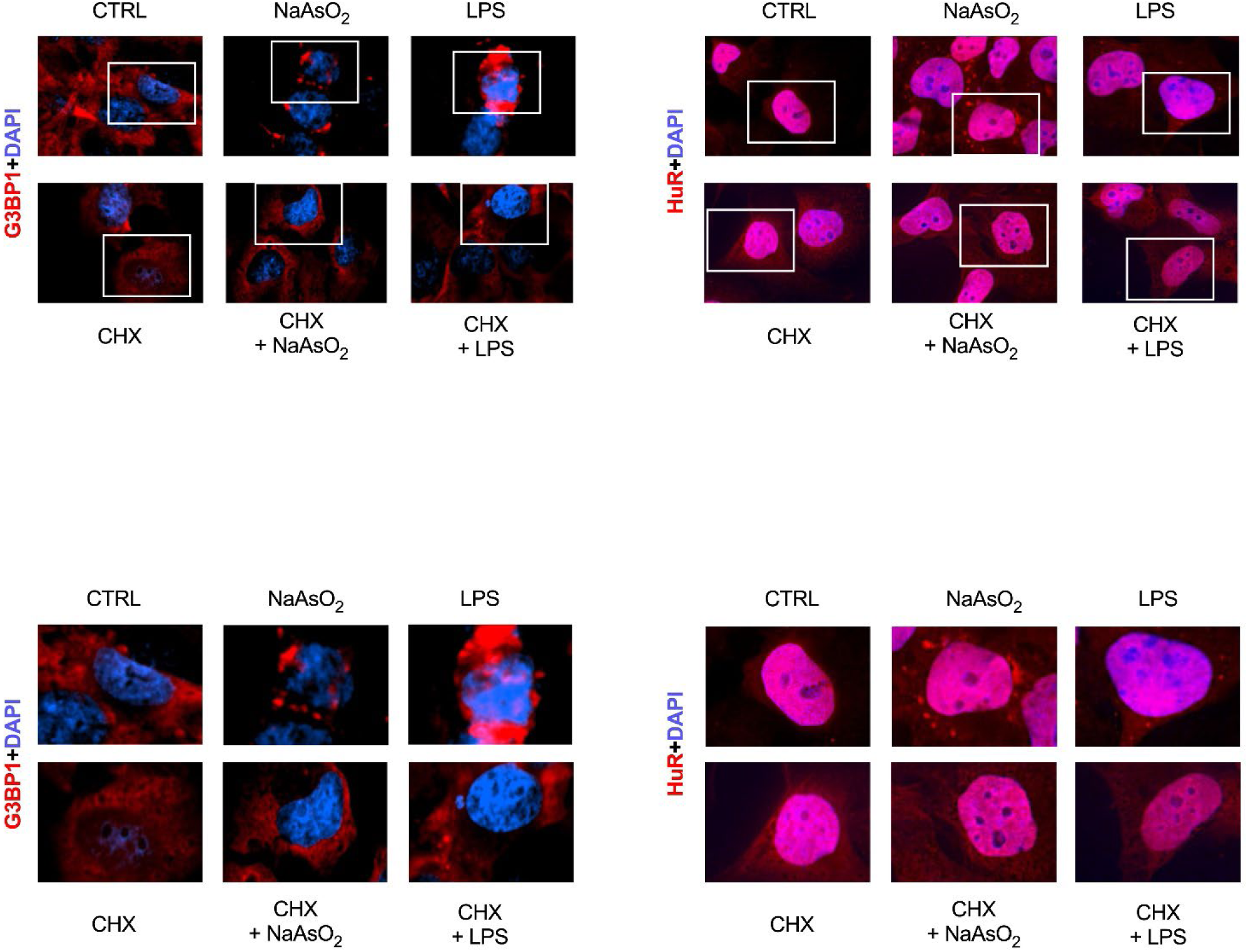
Original immunofluorescence images corresponding to Figure 4D-E. Representative full-field images showing G3BP1 (red) and HuR (red) localization in chondrocytes under the indicated conditions. White boxes indicate the regions selected for magnification and displayed in the main Figure 4D-E. Nuclei were counterstained with DAPI (blue). Images were acquired at 60× magnification.

**Fig. S7.**
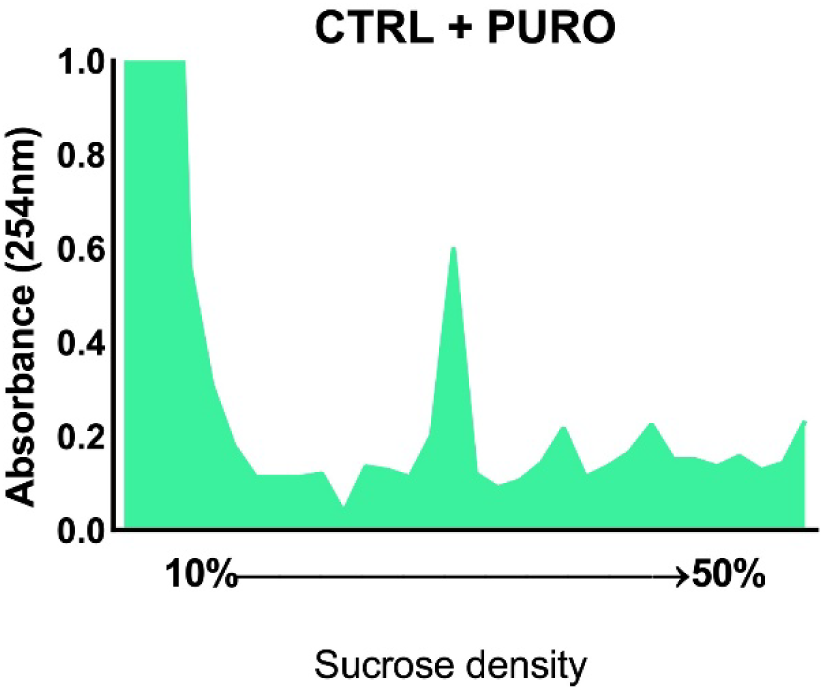
Polysome profiling was performed in C28/I2 chondrocytes and the sample was collected polysome buffer supplemented with puromycin. Identical lysates were loaded onto 10%–50% sucrose gradients, and polysomes were separated by ultracentrifugation. Each fraction was profiled for RNA content. Sucrose-gradient absorbance profile is shown to precisely identify monosome peak.

